# Distinct regulatory mechanisms by the nuclear Argonautes HRDE-1 and NRDE-3 in the soma of *Caenorhabditis elegans*

**DOI:** 10.1101/2024.09.25.615038

**Authors:** Hector Mendoza, Eshna Jash, Michael B. Davis, Rebecca A. Haines, Sarah Van Diepenbos, Györgyi Csankovszki

## Abstract

RNA interference is a conserved silencing mechanism that depends on the generation of small RNA molecules that disrupt synthesis of their corresponding transcripts. Nuclear RNA interference is a unique process that triggers regulation through epigenetic alterations to the genome. This pathway has been extensively characterized in *Caenorhabditis elegans* and involves the nuclear recruitment of H3K9 histone methyltransferases by the Argonautes HRDE-1 and NRDE-3. The coordinate regulation of genetic targets by H3K9 methylation and the nuclear Argonautes is highly complex and has been mainly described based on the small RNA populations that are involved.

Recent studies have also linked the nuclear RNAi pathway to the compaction of the hermaphrodite X chromosomes during dosage compensation, a mechanism that balances genetic differences between the biological sexes by repressing X chromosomes in hermaphrodites. This chromosome-wide process provides an excellent opportunity to further investigate the relationship between H3K9 methylation and the nuclear Argonautes from the perspective of the transcriptome. Our work suggests that the nuclear RNAi and the H3K9 methylation pathways each contribute to the condensation of the X chromosomes during dosage compensation but the consequences on their transcriptional output are minimal. Instead, nuclear RNAi mutants exhibit global transcriptional differences, in which HRDE-1 and NRDE-3 affect expression of their native targets through different modes of regulation and different relationships to H3K9 methylation.

**ARTICLE SUMMARY:** This study examines the transcriptional consequences during the disruption of the nuclear RNAi silencing mechanism in *C. elegans*. Through microscopy and bioinformatic work, we demonstrate that although nuclear RNAi mutants exhibit significantly decondensed X chromosomes, chromosome-wide transcriptional de-repression is not detectable. Downstream analyses further explore the global influence of the nuclear RNAi pathway, indicating that the nuclear Argonautes HRDE-1 and NRDE-3 function through two distinct mechanisms.

## INTRODUCTION

The discovery of RNA interference (RNAi) can be traced back to the climactic days of agricultural biotechnology, right before the first wave of genetically modified produce became available to the public. In an attempt to increase flavonoid content and influence flower pigmentation in petunias transgenically, scientists unexpectedly induced genetic co-suppression, resulting in an overall reduction in pigment production (Napoli et al. 1990; van der Krol et al. 1990). Successive studies determined that this process involved endonucleolytic degradation of RNA (Metzlaff et al. 1997), expanded the finding into other biological systems (Cogoni and Macino 1997; Fire et al. 1998; Pal-Bhadra et al. 1997) and highlighted its potential in a variety of targeted human therapies (Setten et al. 2019).

The mechanism behind RNAi-induced silencing involves the binding of small RNA molecules (sRNAs) by the RNAi-induced silencing complex (RISC), a ribonucleoprotein complex capable of targeted control of gene expression via active degradation of messenger RNA (mRNA) (Bernstein et al. 2001; Hammond et al. 2000), disruption of protein synthesis (Behm-Ansmant et al. 2006a; Behm-Ansmant et al. 2006b; Iwasaki et al. 2009), DNA elimination (Schoeberl et al. 2012) or epigenetic modifications (Verdel et al. 2004; Xie et al. 2004). This last type of RNAi-mediated silencing involves co-transcriptional regulation of targeted genes by the recruitment of histone methyltransferases (HMTs) and the subsequent deposition of histone 3 lysine 9 (H3K9) methylation, a highly conserved marker for heterochromatin formation (Padeken et al. 2022). The overlap between RNAi and chromatin modification allows for a finer genetic control during development.

In *Caenorhabditis elegans*, RNAi-induced transcriptional silencing (RITS) is referred to as nuclear RNAi. Target specificity is assured by the generation of short interfering RNAs (siRNAs) with antisense complementarity to their targets. These RNA strands are 22 nucleotides long and begin with a guanine, hence their categorization as 22G-RNAs (Gu et al. 2009). Worm-specific Argonaute (WAGO) proteins can then associate with and translocate 22G-RNAs to the nucleus, promote heterochromatin formation at specific genomic loci and stall transcript elongation (Buckley et al. 2012; Guang et al. 2010; Guang et al. 2008). This nuclear RNAi silencing pathway recruits the H3K9 HMTs SET- 32, MET-2 and SET-25 and the H3K27 HMT MES-2 (Ashe et al. 2012; Kalinava et al. 2017; Mao et al. 2015; Spracklin et al. 2017). Recruitment of these HMTs to nuclear RNAi targets is dependent on the WAGOs HRDE-1 and NRDE-3, which localize to the nuclei of germline (Buckley et al. 2012; Seroussi et al. 2023) and somatic cells (Guang et al. 2008; Seroussi et al. 2023), respectively.

Most of the published literature on the native targets of the nuclear WAGOs focuses on sequencing of the siRNA populations they bind. Accordingly, HRDE-1 has been associated with silencing of long terminal repeat (LTR) retrotransposons (Ni et al. 2014; Ni et al. 2018), maintenance of the germline (Buckley et al. 2012; Shirayama et al. 2012), propagation of transgenerational silencing (Buckley et al. 2012; Ding et al. 2023; Kim et al. 2021; Rechavi et al. 2014) and interplay with other RNAi silencing pathways (Bagijn et al. 2012; Fischer and Ruvkun 2020; Rogers and Phillips 2020). NRDE-3’s targets include repetitive elements (Padeken et al. 2021; Zhou et al. 2014), ribosomal RNA (rRNA) (Wang et al. 2020; Zhou et al. 2017), intergenic regions (Padeken et al. 2021; Zhou et al. 2014), duplicated genes (Fischer et al. 2011) and have also been linked to the maintenance of heritable silencing (Burton et al. 2011) and to separate RNAi silencing mechanisms (Fischer et al. 2013; Zhuang et al. 2013). The targets of the H3K9 HMTs encompass repetitive elements (McMurchy et al. 2017; Padeken et al. 2019; Zeller et al. 2016) and regulators of tissue differentiation (Methot et al. 2021; Rechtsteiner et al. 2019) and developmental fate (Bian et al. 2020; Gonzalez-Sandoval et al. 2015).

The way in which the nuclear WAGOs and H3K9 methylation coordinate silencing of overlapping genetic targets is complex. For instance, H3K9 methylation is required for silencing of some nuclear WAGO targets in some cases, but not others (Ashe et al. 2012; Kalinava et al. 2017; Lev et al. 2017; Minkina and Hunter 2017; Spracklin et al. 2017). Notably, in a triple *set-32; met-2 set-25* mutant background, most endogenous targets of HRDE-1 are not derepressed (Kalinava et al. 2017; Ni et al. 2014; Ni et al. 2018).

Therefore, H3K9 methylation, specifically H3K9 trimethylation (H3K9me3), cannot be the sole mechanism of silencing directed by HRDE-1. On the other hand, cumulative derepression of LTR retrotransposons in a *hrde-1* mutant background that is also compromised for HMT activity has also been reported (Kalinava et al. 2018; Ni et al. 2018), suggesting that H3K9 methylation can contribute to silencing of nuclear WAGO targets. Previous studies on HRDE-1 focused on germline targets. However, HRDE-1 function clearly impacts somatic development, as exemplified by the derepression of transposons in intestinal cells of *met-2; hrde-1* double mutants (Ni et al. 2018) and the impact on X chromosome decondensation in intestinal cells in a *hrde-1* mutant background (Davis et al. 2022). Understanding the relationship between nuclear WAGOs and H3K9 HMTs provides a great opportunity for the exploration of how these converging silencing pathways can produce specific changes during development.

We previously showed that nuclear WAGOs and H3K9 HMTs also influence packaging of the X chromosomes during dosage compensation (DC) (Davis et al. 2022; Snyder et al. 2016), a mechanism that equalizes gene expression differences between the biological sexes. In *C. elegans*, DC targets both X chromosomes of hermaphrodites (XX), dampening their expression by ∼50% to match that of males (X0) (Meyer 2022). X-specific silencing is promoted by the enrichment of the repressive histone 4 lysine 20 (H4K20) methylation mark, mediated by the histone demethylase DPY-21 (Brejc et al. 2017; Vielle et al. 2012; Wells et al. 2012). Silencing of the hermaphrodite X chromosomes is reinforced by chromatin sequestration and compaction, mediated by the deposition of H3K9 methylation by MET-2, SET-25 and SET-32 (Snyder et al. 2016; Towbin et al. 2012).

DC-defective backgrounds exhibit X chromosome decondensation (Brejc et al. 2017; Davis et al. 2022; Lau et al. 2014; Snyder et al. 2016). Additionally, a wide array of phenotypes and physiological abnormalities have been previously associated with mutations affecting DC, including Dpy, and Egl (for a comprehensive list, refer to the review by Meyer 2022) and can be accompanied by X chromosome derepression (Kramer et al. 2015; Lau et al. 2014; Snyder et al. 2016; Trombley 2024). On the other hand, if ectopically activated in males, DC results in complete lethality due to the inappropriate repression of their single X chromosome (Miller et al. 1988). This male lethality can be rescued by mutations that affect subunits of the dosage compensation complex (DCC) (Miller et al. 1988; Rhind et al. 1995) and, to a lesser extent, by disrupting X chromosome anchoring and compaction via H3K9 methylation (Snyder et al. 2016) and nuclear WAGO function (Davis et al. 2022; Weiser et al. 2017). Rescue by depletion of the H3K9 HMTs or nuclear WAGOs, however, requires partial destabilization of the DC mechanism via a mutation in *sex-1*, a negative regulator of male development in the sex determination pathway of *C. elegans* (Davis et al. 2022; Gladden et al. 2007; Snyder et al. 2016). The potential interplay between DC, H3K9 methylation and the nuclear WAGOs provides a unique perspective to further characterize the nuclear RNAi silencing mechanism.

In this study, we explore the impact of the loss of nuclear RNAi and/or H3K9 methylation on DC and on gene expression in the soma in general. To our knowledge, the impact of the combined loss of nuclear WAGOs and HMTs has only been investigated in the context of silencing of transposable elements by HRDE-1 (Ni et al. 2018). We show that H3K9 and nuclear WAGOs have both overlapping and non-overlapping functions. This finding is based on the additivity resulting from the combined action of the nuclear WAGO and the H3K9 HMTs, reflected as early developmental delays and significantly decondensed X chromosomes during DC.

Analysis of transcriptomic profiles of nuclear RNAi mutants with and without additional mutations in H3K9 HMTs revealed that nuclear RNAi-mediated chromosome compaction does not result in large scale derepression of the X chromosomes. Instead, the absence of HRDE-1, NRDE-3 and the HMTs leads to gene misregulation at the global level. However, H3K9 methylation impacts HRDE-1 and NRDE-3-mediated gene regulation in strikingly different ways. While NRDE-3 and the HMTs exhibit cooperation during gene regulation, the HMTs and HRDE-1 have opposing effects on the regulation of a subset of genes, emphasizing their specific effects in the soma and germline, respectively.

## MATERIALS AND METHODS

### Nematode strains and maintenance

All nematode strains were maintained on nematode growth media (NGM) following standard protocols (Stiernagle 2006). The strains used in this study (Table S1) include N2 (wild-type), CSS419 *set-32 (red11)*; *met-2 (n4256) set-25 (n5021)*, EKM89 *hrde-1 (tm1200)*, CSS415 *set-32 (red11)*; *hrde-1 (tm1200) met-2 (n4256) set-25 (n5021)*, WM156 *nrde-3 (tm1116)* and EKM205 *set-32 (red11)*; *met-2 (n4256) set-25 (n5021); nrde-3 (tm1116)*.

Some strains were provided by the Caenorhabditis Genetics Center (CGC), which is funded by the NIH Office of Research Infrastructure Programs (P40 OD010440).

### Analysis of developmental rate

Synchronized L1 larval worms were obtained by bleaching gravid adults to collect embryos and then allowing them to hatch in M9 buffer (85 mM NaCl, 1 mM MgSO_4_, 22 mM KH_2_PO_4_, 42 mM Na_2_HPO_4_) overnight. L1 animals were then plated on NGM and allowed to grow following standard protocols (Stiernagle 2006) and at 20°C. The number of L4 animals were then manually counted 42 hours post-feeding. Average developmental rates were calculated from multiple biological replicates and compared using ANOVA followed by Tukey’s multiple comparisons performed on GraphPad Prism v10.0 to determine statistical significance, defined as *p* < 0.05.

### RNA extraction, library preparation and bioinformatic analysis

Total RNA was extracted from synchronized L1 worms grown on high growth media (HGM). Synchronized L1 animals were allowed a 3-hour recovery period on NGM with *Escherichia coli* OP50 as food source prior to the extraction procedure. Samples were then lysed by repeated freeze-thaw cycles and subsequent treatment with TRIzol reagent (Invitrogen). Total RNA was isolated using an RNeasy Mini Kit (QIAGEN) with on-column Dnase digestion using RQ1 Rnase-Free Dnase (Promega). Extractions were carried out in three to four biological replicates of each strain. Poly(A)-tailed mRNA enrichment, library preparation and next-generation sequencing were carried out in the Advanced Genomics Core at the University of Michigan. Sample concentration and quality were assessed using a Qubit fluorometer and the TapeStation (Agilent), respectively. Optimal samples (RNA Integrity Number ≥ 7, concentration ≥ 1 ng/μl) were then subjected to poly(A) enrichment using the NEBNext Poly(A) mRNA Magnetic Isolation Module (New England Biolabs). Library preparation was carried out using the NEBNext UltraExpress RNA Library Prep Kit (New England Biolabs). mRNA was fragmented and copied into first strand complementary DNA (cDNA) using reverse transcriptase and random primers. The 3’ ends of the cDNA were then adenylated and adapters were ligated. Products were purified and enriched by PCR for the generation of the final cDNA library. Quality was assessed with a Qubit fluorometer and LabChip (Perkin Elmer). Samples were then pooled and subjected to 151 bp paired-end sequencing using the NovaSeq 6000 S4 Reagent Kit and according to the manufacturer’s instructions (Illumina). BCL Convert Conversion Software v4.0 (Illumina) was used to generate de-multiplexed Fastq files. Reads were trimmed using CutAdapt (v2.3) (Martin 2011) and further evaluated with FastQC (v0.11.8) (Andrews 2010) to determine quality of the data. Reads were mapped to the reference genome Wbcel235 and read counts were generated using Bowtie 2 (v2.4.2) (Langmead and Salzberg 2012) and Htseq 2.0 (v0.13.5) (Putri et al. 2022). Differential gene expression analysis was performed using DESeq2 (v1.42.0) (Love et al. 2014). Differential expression datasets for significantly misregulated genes (*p* < 0.05) with a minimum average of the normalized count values (baseMean > 1) are included in File S2 for reference. Downstream analyses were performed using R scripts and packages and only considered protein-coding transcripts. Gene enrichment analysis was performed using the WormBase gene set enrichment analysis tool with a *q*-value threshold of 0.1 (Angeles-Albores et al. 2018; Angeles-Albores et al. 2016).

### Immunofluorescence

Immunofluorescence staining was carried out following standard protocols (Shaham 2006). Intestinal nuclei from one-day-post L4 animals were dissected on slides in 1X sperm salts (50 mM Pipes pH 7, 25 mM KCl, 1 mM MgSO_4_, 45 mM NaCl and 2 mM CaCl_2_), fixed in 2% paraformaldehyde and frozen on dry ice for 10 minutes. Following three 10-minute washes in PBS with 0.1% Triton X-100 (PBST), slides were incubated with diluted primary antibodies in a humid chamber overnight at room temperature. Slides were then washed three times, for 10 minutes each, with PBST and incubated with secondary antibodies in a humid chamber at 37°C for one hour. Following incubation, the three PBST washes were repeated, with the last wash including the DAPI nuclear stain. Slides were mounted with Vectashield (Vector Labs) prior to storage at -20°C until microscopic examination.

### Fluorescence *in situ* hybridization (FISH) combined with immunofluorescence

Slides of intestinal nuclei from one-day-old L4 animals were prepared as for immunofluorescence up to the first set of PBST washes (see above). Slides were then dehydrated with sequential 2-minute washes in 70%, 80%, 95% and 100% ethanol and allowed to air dry for 5 minutes at room temperature. Detection probes were prepared from degenerate oligonucleotide-primed PCR to amplify yeast artificial chromosomes (YACs) corresponding to sections of the *C. elegans* X chromosome (Lau et al. 2014; Nabeshima et al. 2011; Snyder et al. 2016). Slides were then incubated with 10 μl of the X probe at 95°C for 5 minutes and then at 37°C overnight in a humid chamber. The slides were then sequentially washed at 39°C with 2X saline-sodium citrate (SSC) buffer/50% formamide for 5 minutes (three washes), 2X SSC for 5 minutes (three washes) and 1X SSC for 10 minutes (one wash). The immunofluorescence protocol resumed after these last washes, using antibodies specific for DPY-27.

### Antibodies

Rabbit α-DPY-27 (purified antibody) (Csankovszki et al. 2009) was the only primary antibody used and at a 1:200 dilution. For secondary antibodies, α-rabbit Cy3 (Jackson ImmunoResearch Labs 711-165-152) and α-rabbit FITC (Jackson ImmunoResearch Labs 715-095-152) were used at 1:100 dilutions.

### Microscopy and morphometric analysis

Imaging of intestinal nuclei was achieved with a Hamamatsu ORCA-ER digital camera, mounted on an Olympus BX61 epi-fluorescence microscope with a motorized Z drive. The 60X APO oil immersion objective was used for all images, with Z stacks collected at 0.2 μm increments. All images shown correspond to projection images summed from ∼3 μm. Quantification was performed using the SlideBook 5 software (Intelligent Imaging Solutions, Denver, CO, USA). Segment masks were drawn individually for the DAPI signal (DNA) and the Cy3 signal (X chromosomes) for each nucleus. These masks were established by a user-defined intensity threshold value to exclude background signal and autofluorescence. Nuclear DNA and X chromosome volumes were estimated by assigning a morphometry mask and determining their relative overlap in voxels. The X chromosome territory volume was defined as the ratio of the volume of the Cy3 signal to the volume of the DAPI signal. Averages were then calculated for each mutant background analyzed and one-way ANOVA followed by Tukey’s multiple comparisons was performed on GraphPad Prism v10.0 to determine statistical significance, defined as *p* < 0.05. No more than three nuclei were imaged from any individual animal and identical probe batches were used for each experiment with a wild-type control.

## RESULTS

### The nuclear RNAi and H3K9 methylation machineries modulate development during the larval stages

The loss of H3K9 methylation has been previously shown to delay somatic development, as *met-2 set-25* mutant animals exhibit stochastic growth cycles during the transition from the L1 stage to the L1 stage of the next generation (Zeller et al. 2016). We decided to investigate if the combined action of WAGO and HMTs mutations exacerbated this phenotype given that, although developmentally delayed, *met-2 set-25* mutant animals eventually reach adulthood without any noticeable aberrant morphologies (Zeller et al. 2016). An additional mutation was included in the *met-2 set- 25* background, *set-32*, as only in its simultaneous absence can the H3K9me3 signature be fully abrogated (Kalinava et al. 2018; Kalinava et al. 2017). The strains studied (Table S1) include an “HMTs” mutant (*set-32; met-2 set-25*), two “WAGO” mutants (*hrde-1* and *nrde-3*) and the corresponding “WAGO HMTs” combinations (*set-32; hrde-1 met-2 set-25* and *set-32; met-2 set-25; nrde-3*). We used a modified developmental assay in which synchronized L1 animals were allowed to grow under permissible conditions. The number of animals that reached the L4 stage after 42 hours was then determined to examine if development was occurring at an abnormal rate. In this time frame, most of the wild-type (N2) worms reached the L4 stage (Figure 1, grey bar). In contrast, ∼47% of HMTs mutant animals failed to reach the L4 stage at the expected time (Figure 1, blue bar), consistent with the effect of a *met-2 set-25* mutation (Zeller et al. 2016). Both WAGO mutants exhibited similar developmental delays (∼40% delay for *hrde-1* and 43% delay for *nrde-3*) (Figure 1, green and yellow bars, respectively). Notably, the *hrde-1* HMTs mutant background showed severe developmental delay (∼68%, Figure 1, orange bar), while the effect of the *nrde-3* HMTs combination (∼50%, Figure 1, pink bar) was comparable to that of the HMTs and *nrde-3* mutations alone. These findings corroborate the effect of H3K9 methylation in somatic development and demonstrate the additional involvement of the nuclear RNAi machinery. More importantly, the additivity exhibited in the absence of both HRDE-1 and the HMTs suggests that HRDE-1 contributes to somatic development beyond recruitment of HMTs to its targets.

**Figure 1.**
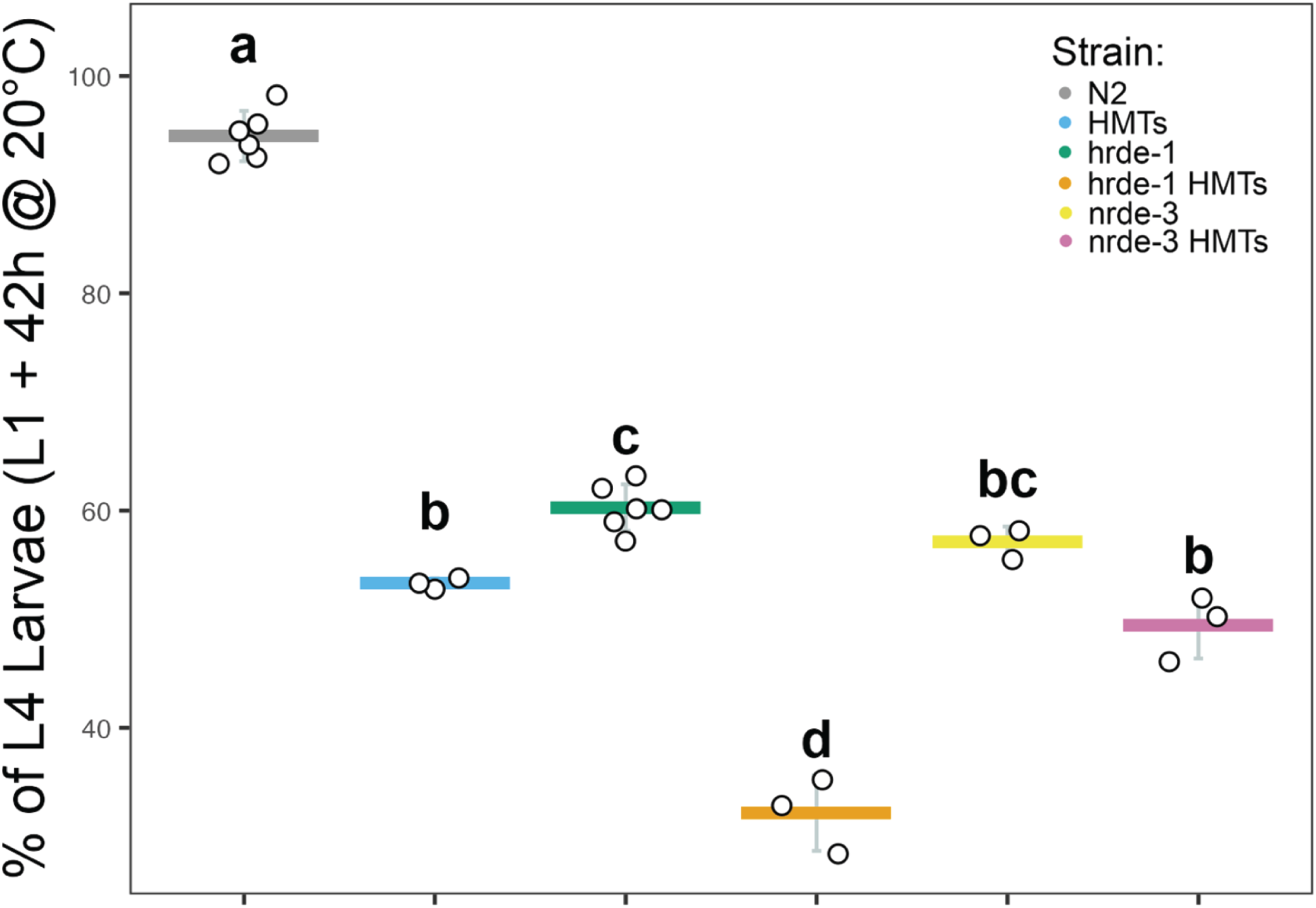
Nuclear RNAi and HMTs mutants display early developmental delay. Percentages of synchronized L1 animals that reached the L4 stage 42 hours post-feeding and grown at 20°C were determined for each mutant strain. Individual points represent biological replicates, with means indicated by horizontal bars; standard deviations, by vertical bars. Different letters indicate statistically significant differences between strains based on one-way ANOVA followed by Tukey’s multiple comparisons (*p* < 0.05).

### HRDE-1 independently contributes to X chromosome compaction during DC

Previous work suggested the involvement of the nuclear WAGOs and H3K9 HMTs in X chromosome repression during in DC (Davis et al. 2022; Weiser et al. 2017), although whether they perform this function together or independently of each other is not known. To determine the impact of the combined loss of HMTs and nuclear WAGOs on X chromosome compaction, the nuclear territories occupied by both X chromosomes in hermaphrodite animals were compared between the different nuclear RNAi and HMTs mutant strains. X chromosome visualization was indirectly achieved by using antibodies against DPY-27, a protein unique to condensin I^DC^, a main component of the DCC (Chuang et al. 1994; Csankovszki et al. 2009). This methodology was validated by X chromosome paint fluorescence *in situ* hybridization (FISH) combined with DPY-27 immunofluorescence, which showed that the DPY-27 signal overlaps with the X chromosome territory in all backgrounds (Figure S1). The corresponding volume of the fluorescent signal was then compared to the total nuclear volume to determine the ratio of the X chromosomes to the entire nuclear DNA volume (Figure S1).

The X chromosome territories in the HMTs, WAGO and corresponding WAGO HMTs mutants were compared in reference to the wild-type background (Figure 2A).

**Figure 2.**
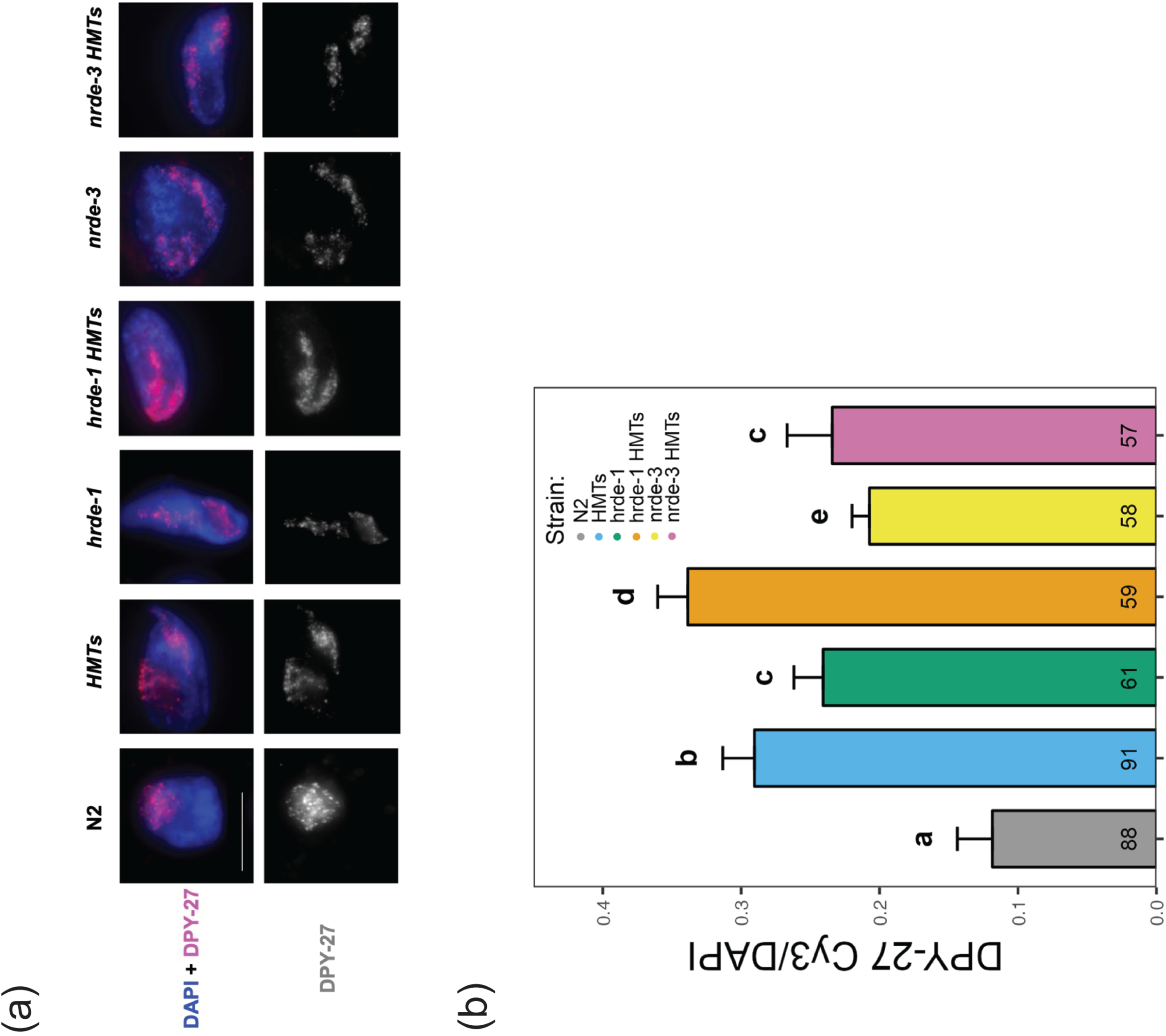
Mutations affecting the nuclear RNAi and H3K9 methylation machineries trigger X chromosome decondensation. **(A)** Intestinal nuclei fixed from one day-old adult (24 hours past L4) animals were subjected to DPY-27 immunofluorescent labelling, using a secondary antibody labeled with Cy3 (red). Labelling of total DNA was achieved using the DAPI nuclear stain (blue). The horizontal white bar represents a 10 μm scale. **(B)** The volume of the X chromosome territory was calculated by determining the ratio of Cy3 to DAPI signal. Bars represent averages of replicates, with standard deviation and number of nuclei analyzed per strain (*n*) indicated above and within bars, respectively. Different letters above bars indicate statistically significant differences between strains based on one-way ANOVA followed by Tukey’s multiple comparisons (*p* < 0.05).

Quantification revealed significantly larger X chromosome territories in the HMTs mutant and in the nuclear WAGO mutants (Figure 2B), consistent with previous studies (Davis et al. 2022; Snyder et al. 2016). Notably, the *hrde-1* HMTs mutant had a larger X chromosome territory compared to both the HMTs and the *hrde-1* mutants (Figure 2B), suggesting an additional role for HRDE-1 in X chromosome compaction separate from the nuclear recruitment of the H3K9 HMTs. This result was not observed for the *nrde-3* HMTs mutant, which had a reduced X chromosome territory compared to the HMTs mutant, but larger than the *nrde-3* mutant, indicating that the impacts on X chromosome packaging for these mutants are not additive (Figure 2B). Instead, the NRDE-3 effect may be primarily attributed to H3K9 methylation and not to WAGO activity alone.

### Nuclear RNAi-mediated X chromosome compaction is decoupled from co- transcriptional control

To assess the impact of nuclear RNAi and H3K9 methylation on X-linked gene expression, we used mRNA-seq to compare global chromosome expression among all autosomes and the X chromosomes in synchronized populations of L1 mutant animals in reference to the wild-type background. We used L1s because we wanted to emphasize somatic tissues, and L1 animals have only two germline precursor cells (Pazdernik and Schedl 2013). Additionally, DC has been shown to begin as early as the ∼40 cell stage, (Dawes et al. 1999), and is well established by the L1 stage (Kramer et al. 2015). mRNA-seq data were first normalized based on read depth and transcript length (FPKM) and only considered protein-coding genes (18141 genes in total). Overall, gene expression at the chromosomal level was mostly unaffected for all mutant strains (Figure 3A-E). Moreover, relative to autosomal expression, the X chromosomes in all mutant strains were not derepressed, suggesting that despite X decondensation, the impact on X-linked gene expression is minimal. Although surprising, this result is consistent with the conditional rescue of inappropriately dosage compensated male animals when the nuclear RNAi mechanism is disrupted (Davis et al. 2022; Weiser et al. 2017) (see discussion). To more specifically look at genes that are sensitive to loss of DC function, an additional dataset was included in the analysis. This data set was generated from synchronized populations of L1 animals treated with *dpy-27* RNAi and consisted of X chromosome targets that were derepressed compared to autosomes in this background (1512 genes in total, “DC Targets”) (Snyder et al. 2016). Based on the overall distribution of the data, X chromosome derepression was still not evident after this filter was applied (Figure 3A-E).

**Figure 3.**
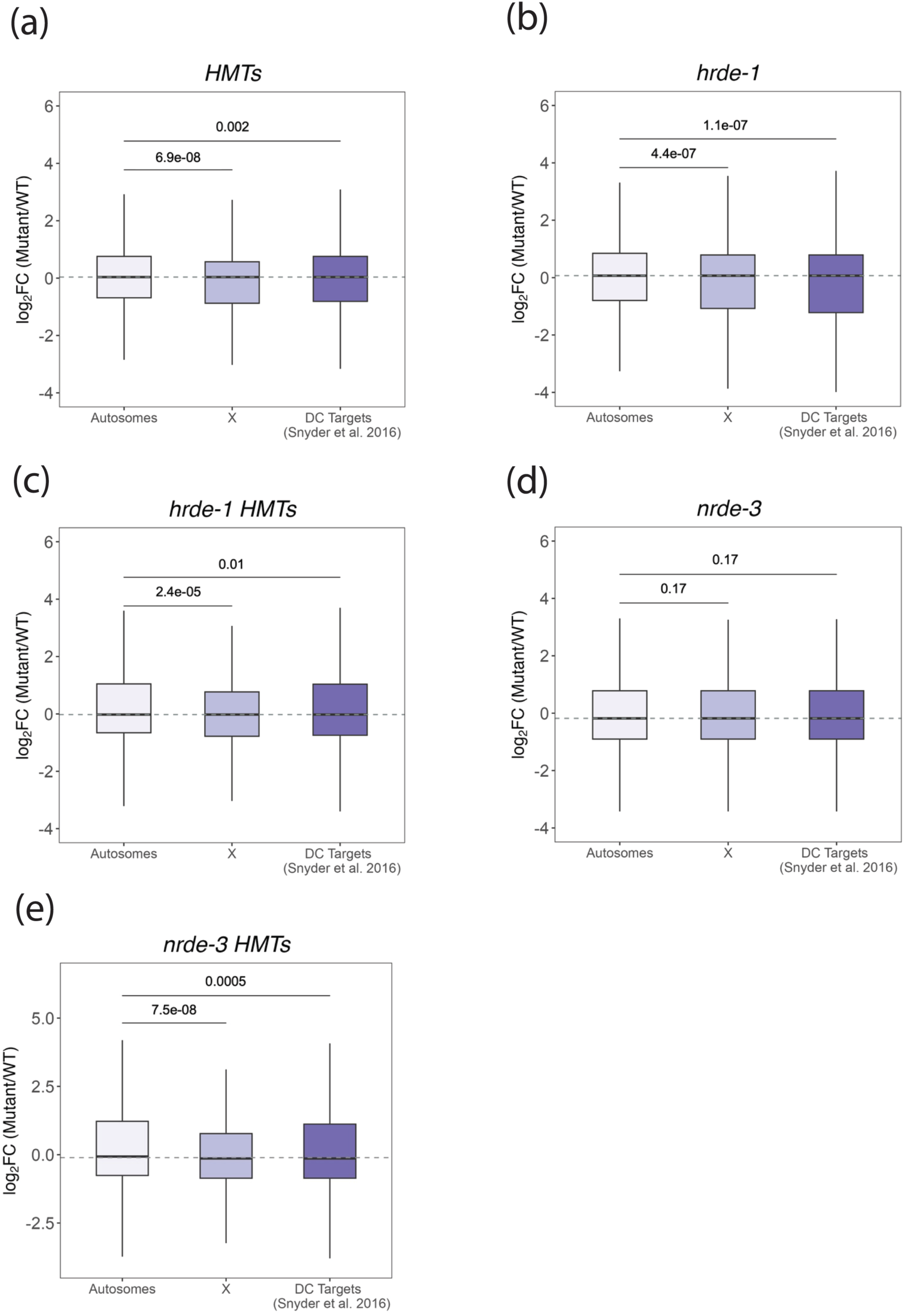
Global X chromosome expression is unaffected by mutations in the nuclear RNAi silencing machinery. Boxplots show the distribution of expression ratios (log_2_ FC) of autosomes, X chromosomes and targets shown to be derepressed (log2FC ≥ 0) in a DC-defective background based on a previous study (Snyder et al. 2016). Medians are indicated by solid black lines and a dashed grey line corresponding to the median of autosomes was included for better visualization of the overall expression shift. The analysis was based on differential expression of each mutant strain relative to the wild- type control: **(A)** HMTs, **(B)** *hrde-1*, **(C)** *hrde-1* HMTs, **(D)** *nrde-3* and **(E)** *nrde-3* HMTs. The Wilcoxon rank-sum test was performed for statistical comparison, with *p*-values indicated above boxplots and statistical significance defined as *p* < 0.05.

### The nuclear RNAi WAGOs and H3K9 HMTs exert global gene expression control

We next assessed the effects of nuclear WAGOs and HMTs on gene expression regulation outside of the context of DC. We first used principal component analysis (PCA) to identify and rank sources of variation among all the strains analyzed. Samples with comprehensively similar transcriptomic profiles should cluster together based on the top two sources of variation (PC1 and PC2). Both WAGO HMTs mutants aggregated with the HMTs mutant, irrespective of the nuclear WAGO involved, indicating that gene expression changes are largely driven by the loss of H3K9 methylation (Figure 4A, “Cluster 1”). The *nrde-3* samples grouped with the wild-type samples (Figure 4A, “Cluster 2”), suggesting that the *nrde-3* mutation has limited impact on protein coding gene expression that more closely resembles the wild-type background. On the other hand, the *hrde-1* biological replicates grouped on their own, suggesting that even though HRDE-1 is a germline expressed gene, it has a significant impact on regulating gene expression in the soma (Figure 4A, “Cluster 3”). This last trend is also in concordance with PCA performed on sRNA sequencing data (Figure 4B) generated in a recent bioinformatic survey of the *C. elegans* AGOs (Seroussi et al. 2023).

**Figure 4.**
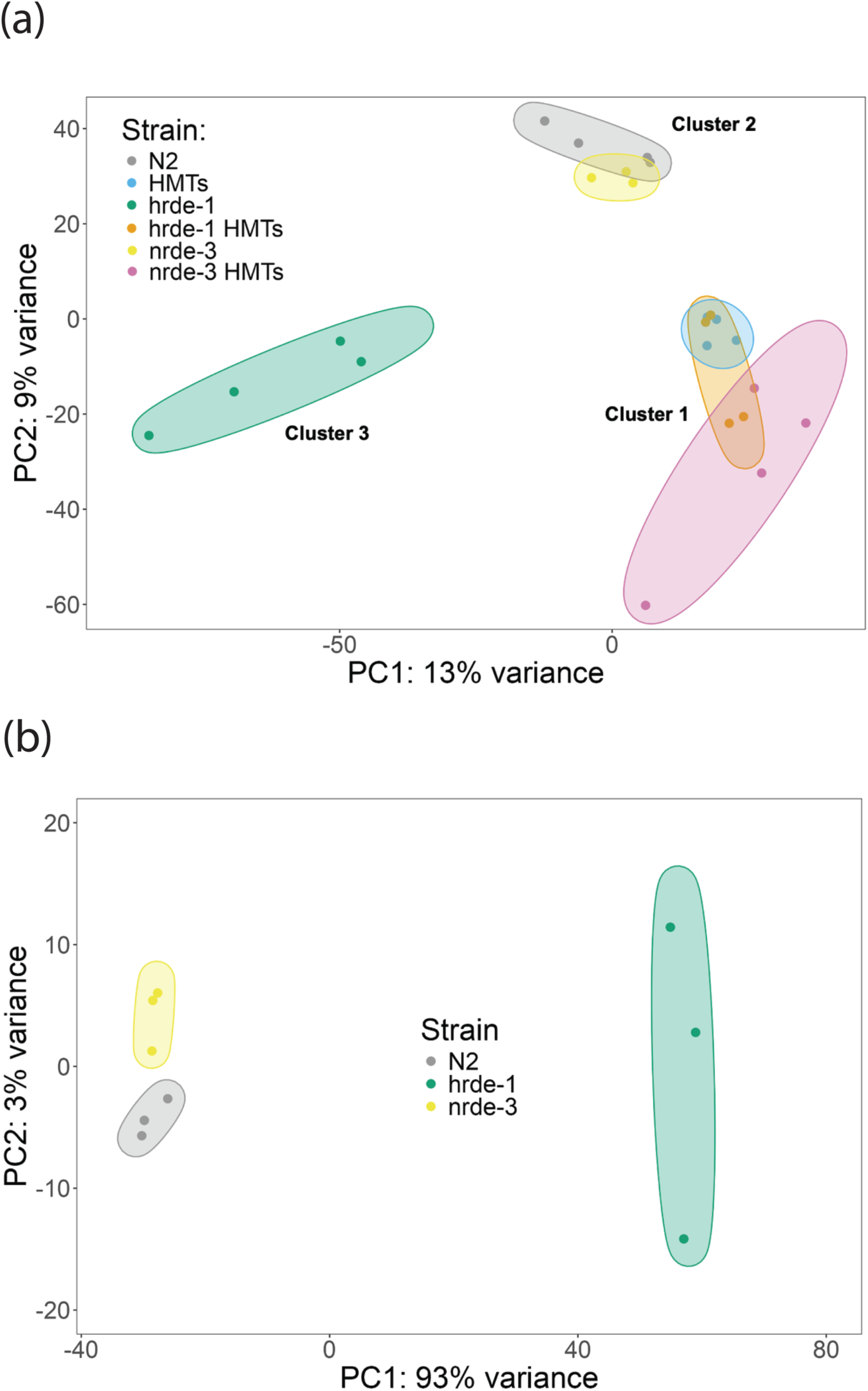
Dimensionality reduction analysis of transcriptomic data reveals distinct roles for HRDE-1 and NRDE-3. **(A)** PCA, based on all *C. elegans* genes, of mRNA-seq data summarizing sample distribution. Individual points represent independent biological replicates for each strain analyzed. **(B)** An additional PCA was performed on sRNA-seq data generated in a separate study (Seroussi et al. 2023)

Previous work suggests that one important role of H3K9 methylation is tissue-specific gene expression (Methot et al. 2021), and specifically repression of germline genes in somatic tissues (Rechtsteiner et al. 2019). To address whether germline-specific genes are overexpressed in the HMTs mutant and to investigate whether nuclear WAGOs contribute to this repression, we examined expression level changes of germline genes. Based on tissue-specific datasets generated in a previous study, we defined “germline genes” as those exhibiting significant upregulation in the young adult gonad compared to the reference background (Spencer et al. 2011). Accordingly, 6052 germline genes were identified in our datasets. To reduce statistical noise, this germline list was trimmed, and we only considered the top 500 germline genes with the highest average expression in the young adult gonad (Spencer et al. 2011). As an additional step, the top 500 germline genes included in the analysis were further examined using WormBase’s enrichment analysis tool (Angeles-Albores et al. 2018; Angeles-Albores et al. 2016). This analysis confirmed that the filtered list corresponded to genes related to germline/reproductive tissue (Figure S2A and File S1).

Relative expression of the top 500 germline genes in each mutant line (mutant/wild-type) was compared to that of all protein-coding genes mapped on the *C. elegans* genome minus the 500 germline genes (17641 “all other genes”). Our analysis revealed that the HMTs have a modest, but statistically significant, overall effect on germline genes, shifting their overall expression above that of all other genes (Figure S2B). Interestingly, disruption of either WAGO alone also led to significant upregulation of germline genes (Figure S2C and S2E). NRDE-3’s influence on germline-specific transcripts is consistent with its ability to translocate to somatic nuclei and trigger the silencing of non-somatic targets (Guang et al. 2008; Seroussi et al. 2023). This finding was also in line with the larval stage chosen for this study (L1), which is predominantly somatic. Since HRDE-1 expression is limited to germline tissues (Buckley et al. 2012; Seroussi et al. 2023), the upregulation detected, albeit much lower than the one caused by NRDE-3, was surprising and likely a secondary consequence of global gene expression changes. A similar trend was observed in the upregulation of germline genes in *hrde-1* HMTs (Figure S2D) and *nrde-*3 HMTs mutants (Figure S2F), although the differences were not statistically significant (*p* > 0.05). We repeated this analysis with other tissue types (intestinal, muscle and panneuronal) and found that their upregulation was not apparent, further supporting the effect specifically on germline-expressed genes (Figure S3).

### The nuclear WAGOs and H3K9 methylation share some native targets

To characterize genes regulated by HMTs, nuclear WAGOs, or both, differential expression analyses for the mutant strains against the wild-type background were filtered, defining statistical significance after multiple testing correction (*padj* < 0.05) and a minimum mean of the normalized count values (baseMean ≥ 1) (Figure 5A-B, colored points). Correlation analyses were then performed between the HMTs mutant and each WAGO mutant. For the *hrde-1*/HMT comparison (Figure 5A), there were 86 and 520 significantly misregulated targets in the HMTs (blue points) and the *hrde-1* (salmon points) mutants, respectively. An additional 40 genes were significantly misregulated in both mutant backgrounds (magenta points). The expression changes of these genes were positively correlated (*R* ∼ 0.7), suggesting that some genes are similarly regulated by HRDE-1 and the H3K9 HMTs. This first set of common targets included 3 anticorrelated targets (Figure 5A, yellow points), highlighting an adversarial relationship between H3K9 methylation and HRDE-1 activity for a limited number of genes.

**Figure 5.**
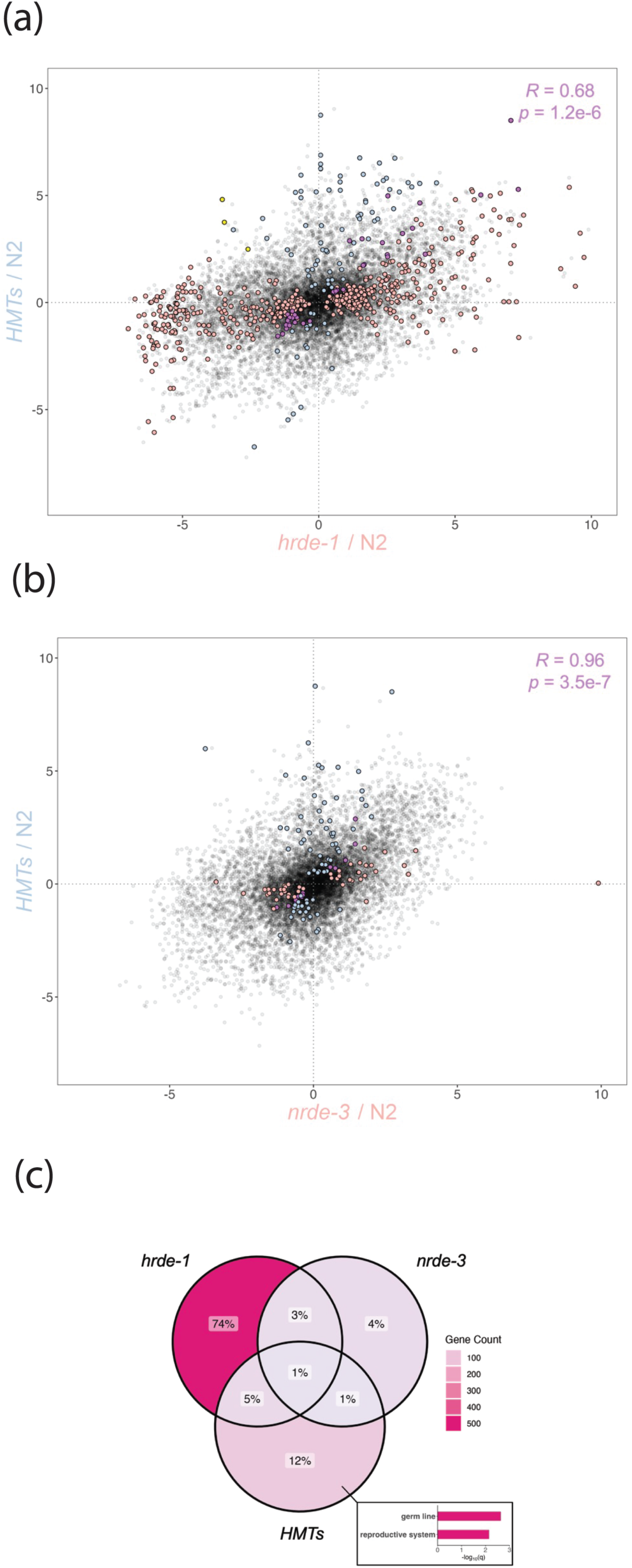
Correlation analysis between the HMTs and single WAGO mutant backgrounds. Comparisons were made between HMTs and **(A)** *hrde-1* or **(B)** *nrde-3* mutants and were based on differential expression analysis of each mutant strain relative to wild-type. Blue and salmon points correspond to significantly misregulated targets (*padj* < 0.05) with a baseMean of at least 1 in the HMTs and each WAGO mutant, respectively. Common targets and the corresponding correlation computation are indicated in magenta. Yellow points indicate anticorrelated common targets. **(C)** Downstream analysis reveals weak overlap between all three datasets examined, with the HMTs influencing a unique gene set significantly enriched for germline tissue (inset).

In the comparison involving NRDE-3 (Figure 5B), the HMTs and *nrde-3* mutations influenced 113 (blue points) and 47 (salmon points) genes on their own, respectively. Only 13 genes were significantly misregulated in both mutant backgrounds (magenta points) and were also strongly positively correlated (*R* ∼ 1). Overlap between all three datasets was notably weak (Figure 5C, ∼1%). Enrichment analysis revealed that the HMTs mutation affected a unique set of genes that were significantly enriched for germline and reproductive tissue (Figure 5C inset and File S1), consistent with our germline validation (Figure S2B) and previous findings (Rechtsteiner et al. 2019). In contrast, although over half of all misregulated genes were affected by the *hrde-1* mutation alone (Figure 5C, ∼74%), enrichment analysis did not produce any significant results. These findings suggest that HRDE-1 affects a heterogeneous gene set that may not be representative of a single specific tissue and/or pathway. The effect of NRDE-3 alone, on the other hand, is less evident, as its absence affects a much smaller gene set (Figure 5C, ∼4%).

### NRDE-3 reinforces H3K9 HMT-mediated silencing

The emergence of significantly misregulated genes in both HMTs and WAGO mutants (Figure 5) suggests potential synergism between H3K9 methylation and nuclear WAGO function. To our knowledge, this phenomenon has only been examined in a single study and only in the context of HRDE-1 and silencing of LTR retrotransposons (Ni et al. 2018). Accordingly, we compared gene expression changes in WAGO HMTs mutants to changes in the HMTs mutant (Figure 6). As expected, a significant portion of genes misregulated in the HMTs background, were also misregulated in the WAGO HMTs backgrounds (Figure 6A and B, magenta points). The expression levels for these genes were strongly positively correlated (R ∼ 1), indicating that for most of these genes, the changes in expression were similar in the mutants and suggesting that these changes were mainly driven by the lack of H3K9 methylation. Consistent with that idea, a large proportion of these genes were misregulated in all three backgrounds (Figure 6C, 24%) and this gene set was significantly enriched for the germline genes (Figure 6C inset and File S1).

**Figure 6.**
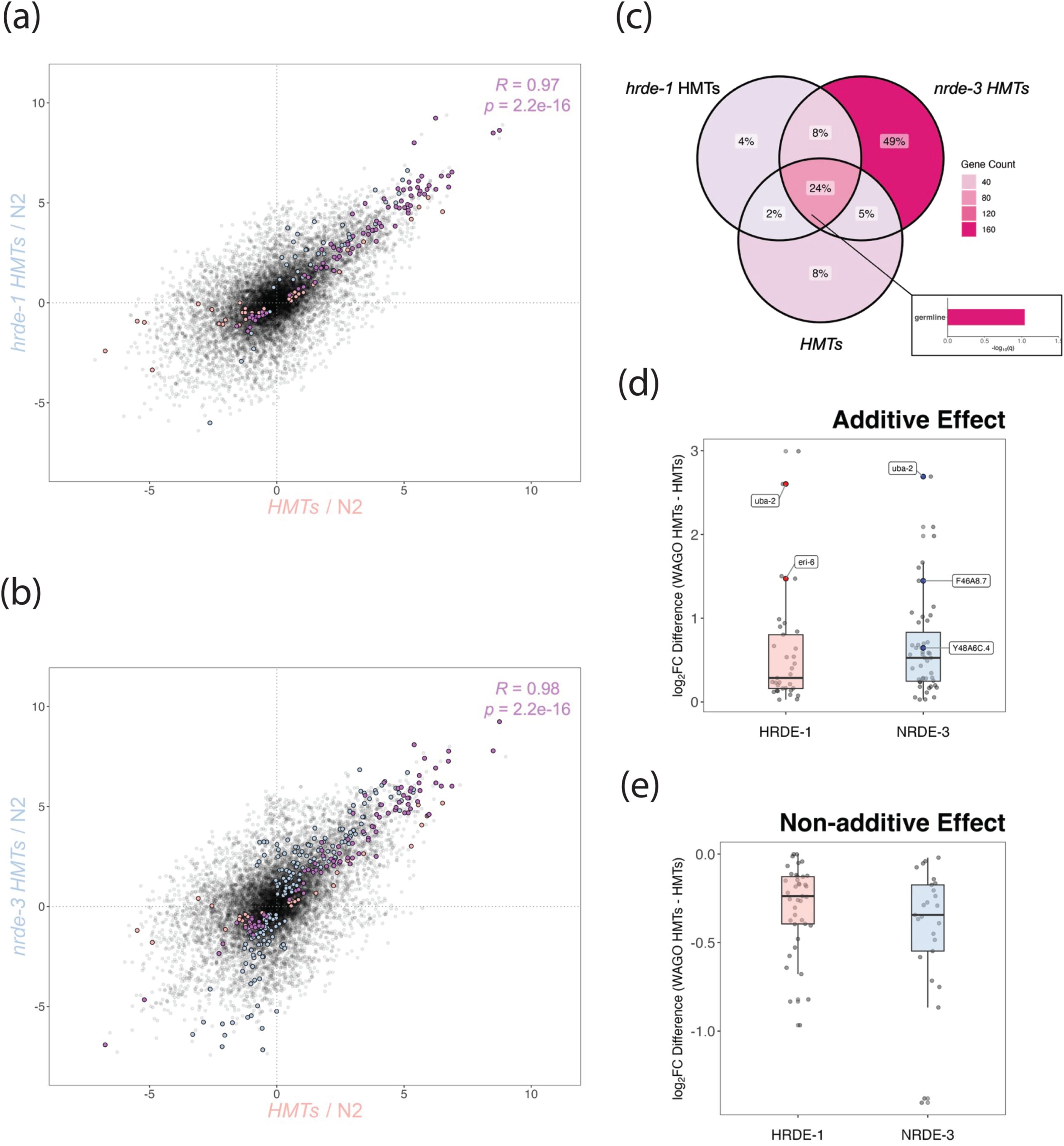
Correlation analysis between the WAGO HMTs and HMTs mutant backgrounds. Comparisons were made between HMTs and **(A)** *hrde-1* HMTs or **(B)** *nrde- 3* HMTs mutants and were based on differential expression analysis of each mutant strain relative to wild-type. Blue and salmon points correspond to significantly misregulated targets (*padj* < 0.05) with a baseMean of at least 1 in the WAGO HMTs and HMTs mutants, respectively. Common targets and the corresponding correlation computation are indicated in magenta. **(C)** Analysis of all three datasets reveals significant overlap and significant germline enrichment (inset, see File S1 for raw data), in addition to a large number of genes only misregulated in the *nrde-3* HMTs background. Differences in relative expression between the upregulated common targets for WAGO HMTs and HMTs mutants were calculated to characterize the effect of WAGO activity on H3K9 methylation. A positive difference (Δlog2FC > 0) was considered additive **(D)**, while a negative difference (Δlog2FC < 0) suggests dampening **(E)**. Boxplots for HRDE-1 (red) and NRDE-3 (blue) targets indicate overall distribution, while individual points are included for the visualization of outliers. Targets exhibiting true additivity or dampening were defined as those with significant misregulation (*padj* < 0.05) in the corresponding differential expression analysis (WAGO HMTs/HMTs), which are labeled and highlighted in red and blue for HRDE-1 and NRDE-3, respectively.

A gene set of considerable size (Figure 6C, ∼49%) was only misregulated in the *nrde-3* HMTs background, suggesting that NRDE-3 and the HMTs redundantly regulate these genes and that their derepression requires the loss of both H3K9 methylation and NRDE-3 function. This is in contrast with the *hrde-1* HMTs mutation (Figure 6C, 4%), accentuating different modes of regulation by the nuclear WAGOs in relation to H3K9 methylation. In these two gene sets, there was no indication for germline bias after enrichment analysis (File S1), reinforcing that germline repression can be predominantly attributed to H3K9 methylation and not WAGO function alone. We then wondered if the genes misregulated in the *nrde-3* HMTs mutant alone (Figure 6C, ∼49%) were misregulated once the effects of H3K9 methylation were removed. For this, we cross-referenced those 160 genes with the differential expression analysis between the *nrde-3* HMTs mutant in reference to the HMTs background (*nrde-3* HMTs/HMTs).

Approximately 17% of those genes were significantly misregulated in this last comparison, indicating that NRDE-3 functions primarily in collaboration with the H3K9 HMTs. For the misregulated genes in the *hrde-1* HMTs mutant alone (Figure 6C, 4%), only 1 of the 12 genes were misregulated in the *hrde-1* HMTs/HMTs comparison. Taken together, these results underscore two distinct regulatory mechanisms, in which NRDE- 3’s transcriptional impact is apparent primarily through collaboration with H3K9 methylation, while HRDE-1 participates in a separate pathway.

Since the HMTs and WAGOs are implicated in gene silencing, we next focused on the genes upregulated in the mutant backgrounds, as these genes have a higher probability of being direct targets. To determine the degree of synergism between nuclear WAGO function and H3K9 methylation, we computed the relative expression (mutant/wild-type) difference (Δlog2FC) of all common targets upregulated in both the WAGO HMTs and HMTs mutants (Figure 6A and 6B, magenta points in the first quadrants).

Differences were then cross-referenced with differentially expressed gene sets generated by comparing the WAGO HMTs mutant against the HMTs background (WAGO HMTs/HMTs). A total of 55 targets were identified as additive, meaning upregulated in the WAGO HMTs compared to HMTs alone (Figure 6D, Δlog2FC > 0), of which only 5 were statistically significant according to the WAGO HMTs/HMTs comparison (red and blue points for HRDE-1 and NRDE-3, respectively). Notably, derepression of *uba-2* was enhanced regardless of the nuclear WAGO involved, and unique targets included *eri-6* for HRDE-1 and *F46A8.7* and *Y48A6C.4* for NRDE-3. A total of 46 genes had dampened repression in the WAGO HMTs compared to the HMTs alone (Figure 6E, Δlog2FC < 0), although none were of statistical relevance based on the corresponding differential expression analysis. Overall, these results suggest that H3K9 methylation and NRDE-3- mediated silencing is redundant for a large number of genes, and additive for a small subset, while HRDE-1’s role is less linked to H3K9 methylation.

### WAGO-mediated silencing can be enhanced and counteracted by H3K9 methylation

To investigate the reverse, whether WAGO-mediated silencing can be reinforced by H3K9 methylation, we carried out correlation analyses between each WAGO HMTs mutant and the corresponding WAGO mutant. Our results indicate that many HRDE-1 targets are not misregulated in the *hrde-1* HMTs mutant (Figure 7A, salmon points), suggesting that the gene expression change caused by lack of HRDE-1 was reversed by the additional loss of the HMTs. We also detected four anticorrelated targets in the context of *hrde-1* HMTs compared to *hrde-1* (Figure 7A, yellow points), three of which were previously detected in the HMTs mutant compared to *hrde-1* (Figure 5A). When the expression patterns of the three anticorrelated targets were compared in the *hrde-1*, HMTs and *hrde-1* HMTs mutant backgrounds, relative to the wild-type control, it is clear that HRDE-1 is having an antagonistic effect to H3K9 methylation (Figure S4). To look for potential redundancy between HRDE-1 and the HMTs, we also examined differential expression of the genes that were only misregulated in the *hrde-1* HMTs mutant alone (Figure 7C, ∼3%) We compared gene expression levels in the *hrde-1* HMTs mutant in reference to the *hrde-1* background (*hrde-1* HMTs/*hrde-1*). Only 9 out of the 20 genes were significantly misregulated in this last comparison. These results again suggest that redundancy with the HMTs is not the main mode of regulation by HRDE-1.

**Figure 7.**
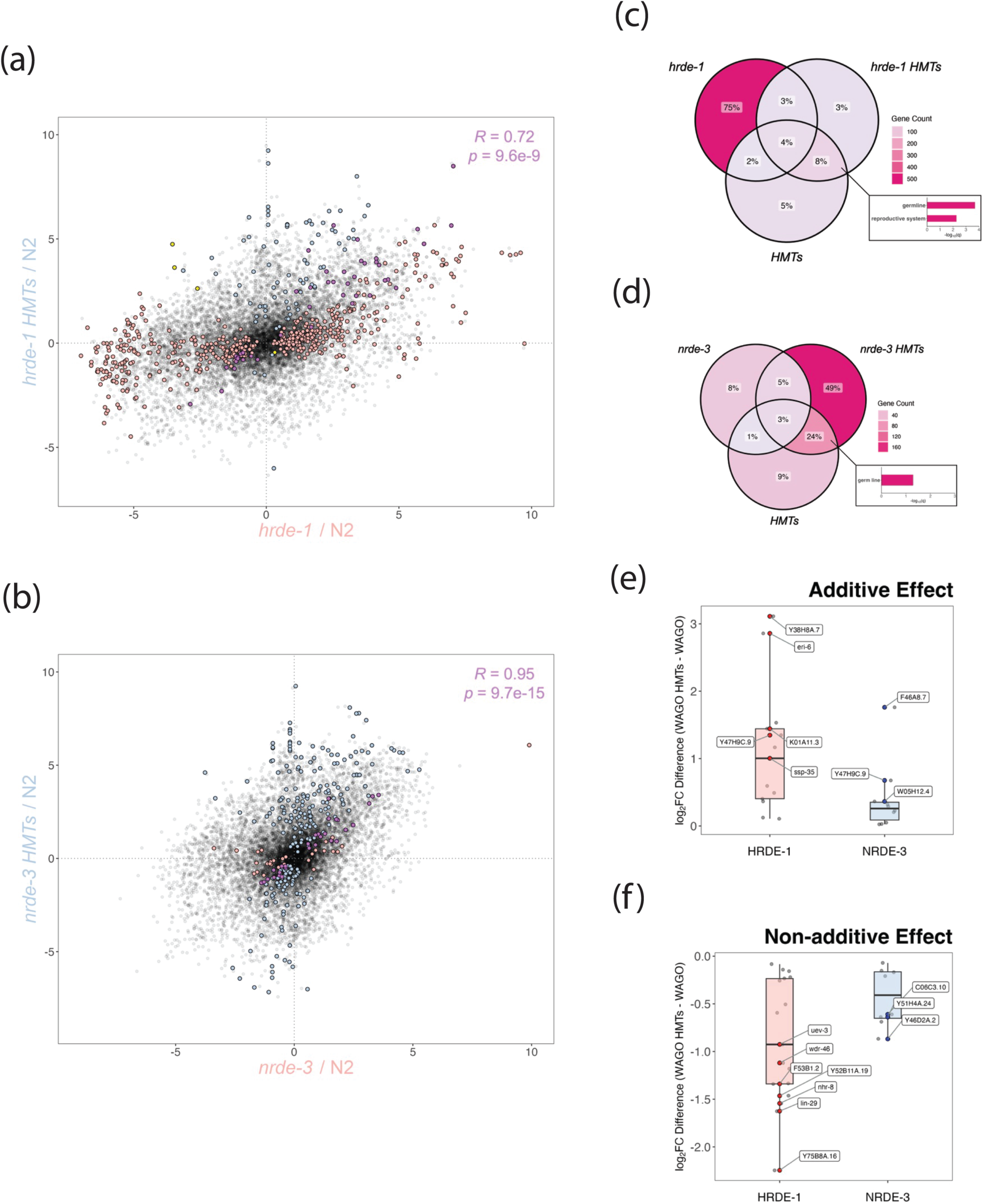
Correlation analysis between the WAGO HMTs and WAGO mutant backgrounds. Comparisons were made between the WAGO HMTs and the single **(A)** *hrde-1* or **(B)** *nrde-3* mutants and were based on differential expression analysis of each strain relative to wild-type. Blue and salmon points correspond to significantly misregulated targets (*padj* < 0.05) with a baseMean of at least 1 in the WAGO HMTs and WAGO mutants, respectively. Common targets and the corresponding correlation computation are indicated in magenta. Yellow points indicate anticorrelated common targets. Overlap analysis between the WAGO, HMTs and WAGO HMTs mutants reveals that **(C)** HRDE-1’s predominant effect involves a mechanism separate from H3K9 methylation, while **(D)** NRDE-3’s effect is primarily dependent on H3K9 HMT activity. Insets illustrate significant enrichment of germline-related terms (see File S1 for raw data). Differences in relative expression between the common targets for WAGO HMTs and WAGO mutants were calculated to characterize the effect of H3K9 methylation on WAGO function. A positive difference (Δlog2FC > 0) was considered additive **(E)**, while a negative difference (Δlog2FC < 0) suggests dampening **(F)**. Boxplots for *hrde-1* (salmon) and *nrde-3* (blue) targets indicate overall distribution, while individual points are included for the visualization of outliers. Targets exhibiting true additivity or non- additivity were defined as those with significant misregulation (*padj* < 0.05) in the corresponding differential expression analysis (WAGO HMTs/WAGO), which are labeled and highlighted in red and blue for HRDE-1 and NRDE-3, respectively.

Contrastingly, NRDE-3’s effect seems to require H3K9 methylation, as very few targets are affected by WAGO activity alone (Figure 7B, salmon points). These differences between NRDE-3 and HRDE-1 were also evident after overlap analysis, with HRDE-1 alone affecting a gene set of considerable size (Figure 7C, 75%) and NRDE-3 affecting more genes only in the context of H3K9 methylation (Figure 7D, 49%). A significant proportion of these 169 genes (∼28%) were also significantly misregulated in the *nrde-3* HMTs/*nrde-3* comparison, reinforcing our previous finding that NRDE-3 and H3K9 methylation enact redundant control on their native targets. Interestingly, germline enrichment was only evident in the common gene sets affected by the WAGO HMTs and HMTs mutant backgrounds for both HRDE-1 (Figure 7C inset and File S1) and NRDE-3 (Figure 7D inset and File S1), with the latter affecting a greater proportion of genes (∼8% vs. ∼24%, respectively). Considering that the germline effect was specific to the HMTs and not the WAGOs based on the previous correlations (Figure 5C), this last finding suggests that the germline-specific effect in somatic tissue is primarily due to H3K9 methylation and not necessarily nuclear WAGO function, which may explain the germline upregulation previously detected in both WAGO HMTs mutants (Figure S2D and S2F).

Lastly, the additivity analyses were repeated for the common upregulated genes between the WAGO HMTs and WAGO mutants (Figure 7A and 7B, magenta points in first quadrant), this time to determine if silencing of WAGO targets is enhanced by H3K9 methylation. To determine statistical relevance, expression differences (Δlog2FC) were then cross-referenced with the WAGO HMTs/WAGO differential expression datasets. A very limited number of genes, a total of 8, exhibited significant additivity (Figure 7E, red and blue points for HRDE-1 and NRDE-3, respectively). The gene *Y47H9C.9*, encoding for a protein of unknown function (Kishore et al. 2020), was influenced by both nuclear WAGOs. HRDE-1 alone influenced 5 genes that included *eri- 6*, which re-emerged from the previous additivity analysis (Figure 6D). In the case of NRDE-3, one of the three additive targets, *F46A8.7*, also appeared in the previous additivity analysis (Figure 6D). Genes exhibiting significant dampened expression (Figure 7F) were WAGO-specific (red and blue points for HRDE-1 and NRDE-3, respectively). Strikingly, our data also indicate an antagonistic relationship between H3K9 methylation and WAGO function, more apparent for HRDE-1 than NRDE-3, suggesting that nuclear WAGOs can target genes outside of the H3K9-mediated silencing pathway.

### HRDE-1 antagonizes H3K9 HMT-mediated repression for a subset of genes

According to the proposed model based on the existing literature, the nuclear WAGOs recruit the H3K9 HMTs to specific gene loci to deposit the heterochromatin marks (Ashe et al. 2012; Guang et al. 2010; Kalinava et al. 2018; Kalinava et al. 2017; Kim et al. 2021; Ni et al. 2014). In our analysis, over 500 genes exhibit significant misregulation in the *hrde-1* mutant, but not in the HMTs mutants (Figures 5A and 5C). Many of these genes remain unaffected in the *hrde-1* HMTs mutant backgrounds (Figure 7) leading us to question the relationship between HRDE-1 and H3K9 methylation for these genes. For this, we performed correlation analysis between *hrde-1*/wild-type and *hrde-1* HMTs/HMTs, with the latter defining the HMTs-independent HRDE-1 effect (Figure 8A). This analysis revealed that the HMTs-independent HRDE-1 effect does not produce significant changes in gene expression (Figure 8A, blue points). This shows that the misregulation of genes in the *hrde-1* mutant background is H3K9-dependent. When we looked at the HRDE-1-independent HMTs effect (*hrde-1* HMTs/*hrde-1*), we noticed that many of the genes that were significantly misregulated in the *hrde-1* mutant were also regulated by the HRDE-1-independent HMTs effect but in the opposite direction (Figure 8B, yellow points, *R ∼* -1). Taken together, these results suggest a model in which the absence of HRDE-1 is causing HMT-mediated silencing of non-specific targets that were not targeted before (Figure 8C and see discussion). This contrasts with what we observe for NRDE-3, where for the majority of genes, significant misregulation is only detected in the absence of both WAGO and HMTs activities.

**Figure 8.**
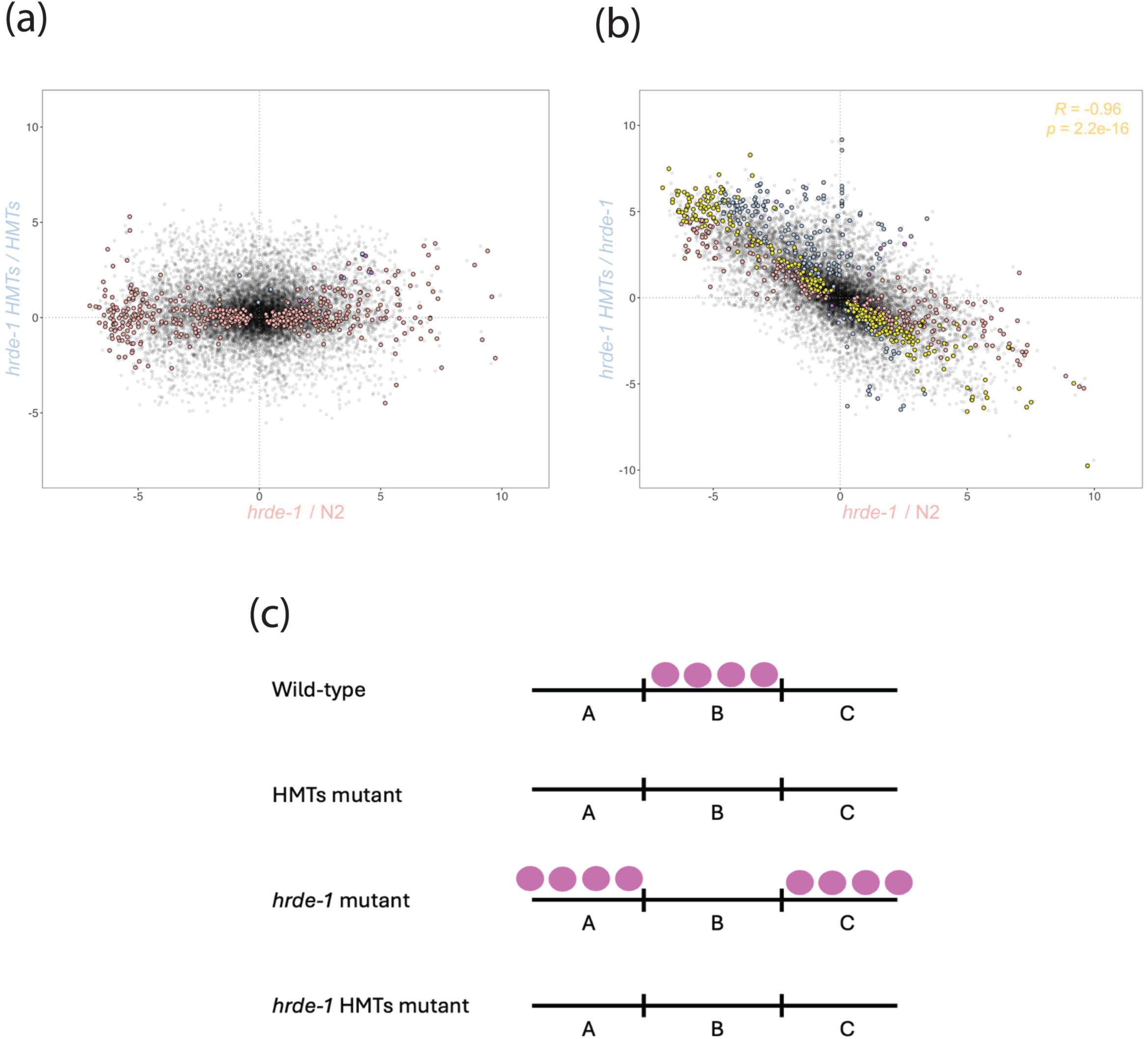
(A) Correlation analysis between the *hrde-1* HMTs/HMTs and the *hrde-1*/wild- type backgrounds further confirms HRDE-1’s unique effect. Blue and salmon points correspond to significantly misregulated targets (*padj* < 0.05) with a baseMean of at least 1 in the *hrde-1* HMTs/HMTs and *hrde-1*/wild-type backgrounds, respectively. Positively correlated targets are indicated in magenta. **(B)** Correlation analysis between the *hrde-1* HMTs/*hrde-1* and *hrde-1*/wild-type backgrounds confirms that HRDE-1 affects a unique set of targets in the opposite direction than H3K9 methylation. Blue and salmon points correspond to significantly misregulated targets (*padj* < 0.05) with a baseMean of at least 1 in the *hrde-1* HMTs/*hrde-1* and *hrde-1*/wild-type backgrounds, respectively. Anticorrelated common targets and the corresponding correlation computation are indicated in yellow, while positively correlated targets are indicated in magenta. **(C)** Our current working model suggests that HRDE-1 drives targeting of H3K9- methylation to specific loci. When HRDE-1 function is lost, H3K9-methylation becomes non-specific and spreads to previously non-methylated genes. H3K9 methylation marks are represented by magenta spheres, while genomic loci are differentiated by different letters.

## DISCUSSION

### Nuclear RNAi as a supporting mechanism for DC

We previously showed that H3K9 methylation and nuclear WAGOs have an impact on X chromosome compaction and DC (Davis et al. 2022; Snyder et al. 2016). In this study, we show that the nuclear RNAi and H3K9 methylation mainly impact DC at the level of X chromatin remodeling (Figure 2), with only minimal changes in overall X-linked gene expression in an otherwise wild-type background (Figure 3). H3K9 methylation is a conserved epigenetic modification found predominantly on silent chromatin (Zhou et al. 2011), although active transcription of genomic regions enriched for H3K9 methylation has been previously reported in *C. elegans* (Kalinava et al. 2017; Methot et al. 2021) and in other model systems (Kim et al. 2007; Vakoc et al. 2005). The limited impact on the transcriptional output of the X chromosomes is consistent with the finding that mutating or depleting the nuclear WAGOs or the H3K9 HMTs in males in which dosage compensation is inappropriately turned on results in rescue, but only in a sensitized background that partially disrupts DC (Davis et al. 2022; Snyder et al. 2016; Weiser et al. 2017). This rescue also requires a mutation in *sex-1*. Accordingly, X-linked gene derepression may be detected in a mutant background in which disruption of DC is achieved by targeting both *sex-1* and HMTs or nuclear WAGOs. H3K9 methylation mediates the packaging of the genome within the nucleus and it works with CEC-4, a nuclear envelope protein that binds to H3K9 methylation marks to anchor the chromosome arms to the nuclear periphery and form active and inactive genome compartments (Bian et al. 2020; Gonzalez-Sandoval et al. 2015). The lack of CEC-4 function alone does not have major impacts on X-linked gene expression, although hermaphrodites do become more sensitive to either the depletion of DPY-27 or a *dpy-21* mutation (Trombley 2024). These observations reveal the involvement of different supplemental mechanisms that support DC, and that loss of any single mechanism may not be enough to significantly disrupt DC and lead to measurable loss of X-linked repression.

### Regulation of germline genes in the soma

Our data also corroborate that H3K9 methylation drives transcriptional control of germline-related genes during early larval development (Figure S2). The most highly expressed germline genes are derepressed to a small degree on all backgrounds tested.

However, enrichment analysis of genes misregulated in the various mutants only showed germline enrichment in the HMTs mutant (Figure 5C) and in the group of genes regulated by both the HMTs and the nuclear WAGOs (Figures 6C, 7C and 7D). These data suggest that the HMTs are more specialized for regulating germline genes in the soma, while the nuclear WAGOs contribute minimally to germline regulation and instead regulate a diverse set of genes on their own.

### The interplay between H3K9 methylation and nuclear WAGO activity

We looked for evidence of synergism between H3K9 methylation and nuclear WAGO function. This analysis allowed us to determine that silencing of H3K9 methylation targets exhibit limited amplification by both nuclear WAGOs (Figure 6). Specifically, we identified four truly additive targets, of which one, *uba-2*, was affected by either nuclear WAGO. This gene encodes for one of the heterodimeric subunits of a ubiquitin-activating enzyme (E1) involved in the charging of SMO-1, the only small ubiquitin like modifier (SUMO) found in *C. elegans* (Choudhury and Li 1997; Surana et al. 2017).

Previous RNAi experiments showed that depletion of SMO-1 leads to embryonic arrest (Fraser et al. 2000). Subsequent studies demonstrated that the process of SUMOylation itself is an essential post-translational modification during cellular division and cell cycle progression (Pelisch et al. 2014; Tsur et al. 2015) in a variety of tissues that include the germline (Kaminsky et al. 2009). This finding could explain the early developmental delay previously detected in all of our nuclear RNAi mutants (Figure 1).

*eri-6* exhibited augmented silencing in the context of HRDE-1. This gene encodes for a component of the enhanced (Eri) RNAi pathway (Fischer et al. 2008), suggesting biological crosstalk between two separate RNAi silencing mechanisms. This observation is also consistent with other findings, in which disruption of HRDE-1 was associated with abnormal mRNA levels of *eri-6* (Ni et al. 2014; Rogers and Phillips 2020). In contrast, NRDE-3 enhanced silencing of *F46A8.7*, enriched in neural tissue and involved in *daf-2*-dependent longevity (Kishore et al. 2020; Ruzanov et al. 2007), and *Y48A6C.4*, encoding a putative component of the MML1 complex, which is crucial during larval development (Kishore et al. 2020).

We then wondered if nuclear WAGO targets also experience additive silencing by H3K9 methylation (Figure 7). A total of seven genes exhibited amplified silencing, with *Y47H9C.9* influenced by either nuclear WAGO. *eri*-6 reappeared in the context of HRDE- 1 only, along with three other genes: *Y38H8A.7* and *K01A11.3*, both encoding uncharacterized proteins (Kishore et al. 2020); and *ssp-35,* involved in sperm development (Kishore et al. 2020). For NRDE-3, genes exhibiting enhanced silencing included *W05H12.4*, encoding for a protein involved in embryonic development (Kishore et al. 2020); and *F46A8.7*, which also appeared in the previous additivity analysis (Figure 6).

### Distinct regulatory mechanisms by the nuclear Argonautes and H3K9 methylation

While synergism between nuclear WAGO action and HMTs was only seen for a limited number of genes, our global transcriptomic analyses revealed fundamentally different mechanisms of regulation for a larger set of genes. The majority of the genes regulated by NRDE-3 are redundantly regulated by NRDE-3 and H3K9 methylation. These genes are only misregulated in worms lacking both the HMTs and NRDE-3 (Figure 6C and Figure 7D). By contrast, very few genes are regulated redundantly by the HMTs and HRDE-1. Instead, the majority of HRDE-1 regulated genes are only misregulated in the *hrde-1* mutant background, and for many their expression does not change in the HTMs background. Furthermore, many of these genes are no longer misregulated in the *hrde-1* HMTs background, indicating an antagonistic effect between HRDE-1 and H3K9 methylation. These results are different from the mechanism behind nuclear regulation of non-coding targets like the Cer3 and Cer8 retrotransposons in the germline. For these targets, HRDE-1 function is essential, while H3K9 methylation is dispensable but can synergistically function with HRDE-1 to significantly increase the level of misregulation (Ni et al. 2018). The differences imply either a fundamental difference between the regulation of protein coding genes and regulation of transposons, or a difference between the impact of HRDE-1’s function in the soma versus the germline. We suggest a model in which HRDE-1 function is essential to guide H3K9 methylation in a locus-specific manner and in its absence, the heterochromatin marks can be mistargeted. The spread of non-specific heterochromatin marks and the fact that this was not the case for NRDE-3 may explain why at the global level the absence of HRDE-1 alone separates itself from both the wild-type and HMTs backgrounds (Figure 4). HMTs/WAGO (Sheet 3).

## DATA AVAILABILITY

RNA-seq datasets generated in this study are available at the NCBI GEO database, accession GSE277005. ncRNA-seq datasets previously published and used in this study are available at the NCBI GEO database, accession GSE208702.

---

Figure S1. Maximum intensity projections of three-dimensional image stacks of X chromosome territories. **(A)** Intestinal nuclei fixed from one-day-old adult animals were subjected to X chromosome paint fluorescence *in situ* hybridization (Cy3 labelled probes, red). Subsequent immunofluorescent labelling was carried out using α-DPY-27 antibodies and FITC-conjugated secondary antibodies (green). Labelling of total DNA was achieved using the DAPI nuclear stain (blue). The horizontal white bar represents a 10 μm scale. **(B)** Nuclear DNA, DPY-27 and X chromosome territories were identified by assigning a corresponding morphometry mask (voxels) and determining relative overlap.

Figure S2. Repression of germline-specific transcripts in the soma is an HMT-dependent process reinforced by nuclear WAGOs. Germline genes were defined as targets with significant upregulation in the young adult gonad in reference to the background control and as described in a previous study (Spencer et al. 2011). An additional threshold was applied to the analysis by reducing the germline list to the top 500 genes with highest average expression. **(A)** The filtered list was significantly enriched for germline-related terms based on enrichment analysis (for raw data, see File S1). These genes were subsequently identified in the datasets generated for each nuclear RNAi mutant line in reference to the wild-type background. Boxplots correspond to average expression of top 500 germline (red) and remaining (blue) genes, with medians indicated by solid black lines. A dashed grey line corresponding to the median of the non-germline targets was included for better visualization of the overall expression shift. The Wilcoxon rank-sum test was performed for statistical comparison, with *p*- values indicated above boxplots and statistical significance defined as *p* < 0.05. Panels correspond to the different mutant backgrounds examined: **(B)** HMTs, **(C)** *hrde-1*, **(D)** *hrde-1* HMTs, **(E)** *nrde-3* and **(F)** *nrde-3* HMTs.

Figure S3. Non-germline genes do not exhibit significant upregulation in the absence of the nuclear RNAi machinery or the HMTs. Intestinal, muscle and panneuronal genes from larval samples were defined as targets with significant upregulation in reference to their corresponding background control and as described in a previous study (Spencer et al. 2011). The lists consisted of 6592, 5943 and 5930 intestinal, muscle and panneuronal targets, respectively. Boxplots correspond to average gene expression in the specific tissue (red) and remaining (blue) genes, with medians indicated by solid black lines. A dashed grey line corresponding to the median of the non-germline targets was included for better visualization of the overall expression shift. The Wilcoxon rank-sum test was performed for statistical comparison, with *p*-values indicated above boxplots and statistical significance defined as *p* < 0.05. Panels correspond to the different mutant backgrounds examined: **(A-C)** HMTs, **(D-F)** *hrde-1*, **(G-I)** *hrde-1* HMTs, **(J-L)** *nrde-3* and **(M-O)** *nrde-3* HMTs.

Figure S4. HRDE-1 has an antagonistic effect to H3K9 methylation on three specific genes (anticorrelated points). Vertical axis corresponds to differential expression of the mutant background relative to the wild-type control. The analysis considered the *hrde-1*, *HMTs* and *hrde-1* HMTs mutant backgrounds.

**File S1.** Raw data for enrichment analyses related to Figures S2A (Sheet 1), 5C (Sheet 2), 6C (Sheet 3) and 6C + D (Sheet 4). Analyses were performed using the WormBase gene set enrichment analysis tool with a *q*-value threshold of 0.1 (https://wormbase.org/tools/enrichment/tea/tea.cgi).

**File S2.** Differential expression datasets filtered according to statistical significance (*p* < 0.05) and the average of the normalized count values (baseMean > 1). The comparisons include mutant/wild-type (Sheet 1), WAGO HMTs/HMTs (Sheet 2) and WAGO

